# Risk management strategies in social grooming among Tonkean and rhesus macaques

**DOI:** 10.64898/2026.10.13.757831

**Authors:** Mathieu Legrand, Sarah Silvère, Natacha Bande, Fannie Beurrier, Gaia Poincelet, Marc Joosten, Adam Provin, Hélène Meunier, Sébastien Ballesta

**Affiliations:** Centre de Primatologie de l’Université de Strasbourg, Niederhausbergen, France; Laboratoire de Neurosciences Cognitives et Adaptatives, UMR 7364, Strasbourg, France

**Keywords:** Non-human primates, conflict management, allo-grooming, eye-contact, social tolerance

## Abstract

Group-living individuals interact constantly, maintaining cohesion, stability, and security within their social groups. However, these interactions can entail risks that individuals must manage to optimize their social navigation through complex social groups, such as those of non-human primates. Here, we investigated how macaques manage risk during social grooming by analysing its somatic modalities, including body orientation and body site being groomed. We compared two macaque species with contrasting social styles: the more despotic rhesus macaque and the more tolerant Tonkean macaque. Using dyadic analyses including species, kinship, age difference, rank difference, and male presence as predictors, we found marked species differences in grooming patterns. Tonkean macaques engaged more frequently in face-to-face grooming and less in face-to-back grooming compared to rhesus macaques. They also groomed body sites associated with high eye-contact probability, such as the head, neck, and chest, more often than rhesus macaques. Kin dyads groomed these body sites more often, particularly in rhesus macaques. These findings suggest that avoiding body sites associated with potential eye-contact may constitute a risk management strategy during grooming. Overall, our results reveal a previously overlooked dimension of primate social cognition that may shape dyadic relationships and social network structure.

## Introduction

Group living requires a balance between the benefits it provides, such as improved protection from predators and greater access to food, and the costs it entails, including the spread of disease and competition for resources (Balasubramaniam *et al*., 2014; Majolo *et al*., 2008; Kappeler *et al*., 2015; Markham & Gesquiere, 2017). In non-human primates (NHPs), social relationships represent long-term investments that can be jeopardized by conflicts of interest, especially when disputes escalate into damaging fights (Kutsukake & Castles, 2001). In fact, risk and uncertainty are inherent to primates’ social interactions (Bohnet *et al*., 2008; Calcutt *et al*., 2019). These exchanges can escalate into agonistic interactions, the costs of which arise from injury (Massey *et al*., 2022), energetic expenditure (de Waal & Davis, 2003) and damage to the relationships (Cords & Aureli, 2000). Maintaining social bonds therefore depends on adaptive behavioural mechanisms observed in both humans and NHPs (Cords & Killen, 1998). The primatological literature has extensively reported on conflict management strategies, such as avoidance, submission patterns and appeasement signals, as well as post-conflict management behaviours such as reconciliation (Cords & Aureli, 2000; Palagi *et al.,* 2008; Ballesta & Meunier, 2023). By contrast, little is known about the behavioural strategies employed during social interactions to reduce the likelihood or cost of social conflict. Identifying such strategies is also key to understanding how primates manage conflict effectively.

Primates can use two broad types of strategies to minimize risks during or before social interactions. The first involves social association strategies, in which individuals adjust the distance, the amount of time spent in interaction or selectively choose partners (Silk *et al*., 1981; Furuichi, 1983; De Waal & Luttrell, 1986; Borries, 1992; Butovskaya *et al*., 1994; Kapsalis and Berman, 1996; Birky & Su, 2005; Roubová *et al*., 2015; Balasubramaniam *et al*., 2017; Kaufhold et al., 2025). Social association strategies allow individuals to prioritize interactions with preferred partners and reduce the risks of conflict (Smuts & Watanabe, 1990). Spatial proximity is strongly influenced by socio-demographic factors: related individuals (Furuichi, 1983), those of similar rank or age, and those of the same sex are often more likely to associate closely (De Waal & Luttrell, 1986; Cords, 2002; Birky & Su, 2005). While selective spatial associations can lower conflict risk, they cannot eliminate it entirely.

Close interactions remain essential for maintaining group cohesion, meaning that individuals must also develop behavioural mechanisms to limit the risk of injury during such interactions. These risks are particularly relevant during behaviours involving close physical contact, such as social grooming. The second strategy involves behavioural adaptations, such as modifying body positions or grooming different body sites (hereafter ’grooming somatic modalities’), to reduce vulnerability during close contact and thereby minimize the potential costs of social interactions (Boccia, 1989; Boccia *et al*., 1989; Barton, 1985; Thierry *et al*., 1990; Moser *et al*., 1991; Allanic *et al*., 2020; Allanic *et al*., 2021; Pfoh *et al*., 2021). These behavioural strategies are expected to vary according to socio-demographic factors such as kinship, sex, age, dominance hierarchy (Moser *et al*., 1991; Borries, 1992; Allanic, *et al*., 2021), and the overall social tolerance characteristic of the species (Boccia, 1989; Thierry, 2000, 2007).

Grooming involves one individual manipulating the fur of another using the hands or mouth (Hutchins & Barash 1976; Boccia, 1983; Barton, 1985). It is a fundamental behaviour for maintaining group stability and cohesion (Dunbar, 1991) and can account for up to 17% of NHPs’ daily activity budget (Lehmann *et al*., 2007). Beyond its hygienic function (Hutchins & Barash, 1976; Tanaka & Takefushi, 1993; Zamma, 2002), grooming plays a crucial social role by strengthening bonds (Cheney, 1992; Dunbar, 2010), reducing tension (Keverne, *et al*., 1989; Aureli & Yates, 2010; Grandi & Ishida, 2015), and facilitating access to social resources such as sexual partners, food, or infants (Gumert, 2007a, Gumert, 2007b). Grooming is not randomly distributed within groups: individuals tend to groom kin (“kin-biased attractiveness”, Seyfarth, 1977) and higher-ranking individuals (“rank-related attractiveness”, Kapsalis & Berman, 1996; Balasubramaniam & Berman, 2017). Consequently, the choice of grooming partners reflects a combination of social preferences and strategic behaviour influenced by socio-demographic factors (Boccia 1983, Schino, 2001, Roubová *et al*, 2015).

Despite its advantages, close proximity during grooming can increase the likelihood of conflict (Schaffner & Aureli, 2005). Body positions adopted during grooming may expose particularly vulnerable areas, increasing potential injury risk if aggression occurs (**Figure 1)**. However, only a small fraction of the literature has considered relative body orientation (e.g., face-to-face, face-to-back; Allanic *et al*., 2021; Thierry *et al*., 1990) or the specific body sites groomed (e.g., neck, chest, back, belly; Barton, 1985; Moser *et al*., 1991). The influence of socio-demographic factors on grooming somatic modalities remains poorly understood, with studies reporting inconsistent results across species (Moser *et al*., 1991; Pfoh *et al*., 2021). Nevertheless, evidence suggests that social affinity (Pfoh *et al*., 2021), sex (Thierry *et al*., 1990), and dominance rank (Moser *et al*., 1991) affect grooming somatic modalities in ways consistent with risk management strategies. Monkeys may protect vulnerable body sites by avoiding positions that expose vital areas (Barton, 1985; Pfoh *et al*., 2021), particularly in males (Moser *et al*., 1991). Although the relative vulnerability of body sites has not been systematically quantified in NHPs, it is plausible that primates adopt grooming postures that reduce perceived risk.

**Figure 1.**
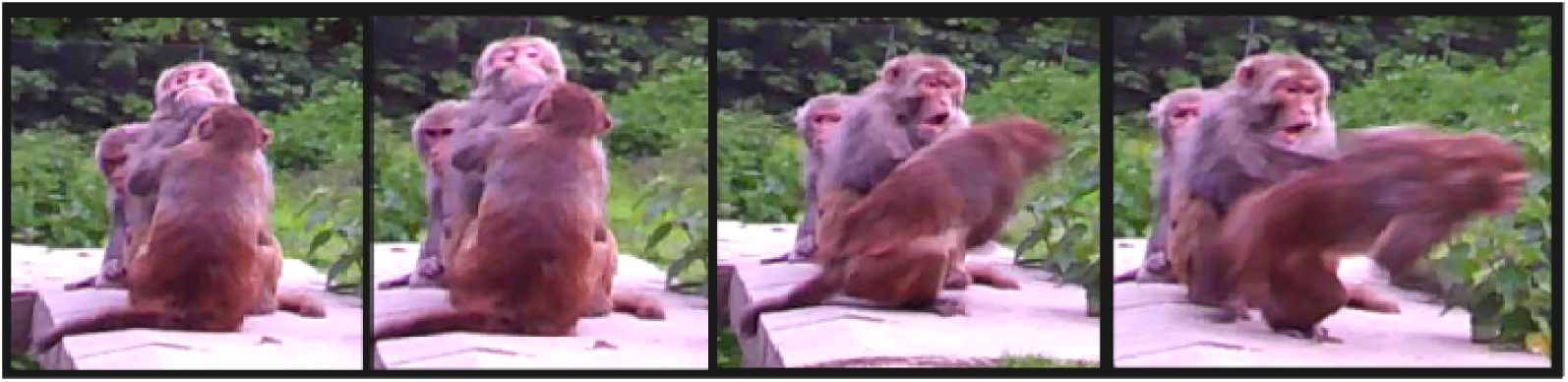
Example of a social grooming session escalating into a physical aggression in rhesus macaque (*Macaca mulatta*). In this sequence, the alpha male was groomed on the neck, in face-to-face position, by a young female. After the two conspecifics made eye-contact (second panel), the dominant male displayed an open-mouth threat, and the female fled before being attacked by her former grooming partner. These events last around two seconds and illustrate the risk associated with social grooming interactions, particularly when individuals are facing each other.

Another potential strategy for minimizing conflict is to avoid eye-contact during grooming. In primates, direct gaze is a powerful social signal that can convey both affiliative and agonistic intent (Emery, 2000; Ballesta & Duhamel, 2015). Eye-contact is frequently used as a threat display, especially among Afro-Eurasian monkeys, where dominant individuals typically initiate it (Harrod *et al*., 2020). It has been proposed that macaques adopt grooming positions that reduce the likelihood of mutual gaze when the risk of conflict is high (Allanic *et al*., 2020; Moser *et al*., 1991). However, this hypothesis has not been directly tested, as the probability of eye-contact according to body orientation or grooming target has never been objectively measured. Tolerance for eye-contact varies across species, depending on social style: more despotic species are more likely to use gaze as a threat (Harrod *et al*., 2020). Consequently, the present study hypothesized that eye-contact may carry a higher risk of aggression in socially intolerant species, influencing how individuals position themselves during grooming.

Macaques offer an excellent model for studying these processes because they display a wide range of social tolerance, classified into four grades from highly despotic (grade 1, e.g., *Macaca mulatta*) to highly tolerant (grade 4, e.g., *Macaca tonkeana*) (Thierry, 2000, 2007, 2022; DeTroy *et al*., 2022). In despotic species, reconciliation rates are lower, conflicts are more severe, and social relationships are marked by nepotism and strong asymmetries (Demaria, & Thierry, 2001). These interspecific differences in social tolerance may thus predict differences in how individuals manage risks during grooming interactions. Specifically, despotic species may be expected to adopt more risk-avoidant grooming postures, minimizing eye-contact and avoiding exposure of vulnerable body regions, than tolerant species.

Addressing the above gaps, the present study investigates how species-typical social style and socio-demographic factors influence grooming somatic modalities. We focus on two macaque species with contrasting social tolerance grades: the despotic rhesus macaque (*M. mulatta*, grade 1) and the tolerant Tonkean macaque (*M. tonkeana*, grade 4). We further introduce an objective metric, the *Eye-Contact Probability* (ECP), to quantify the likelihood of eye-contact during grooming according to relative body orientation and groomed body sites. We hypothesize that despotic species, characterized by low social tolerance, will adopt grooming configurations minimizing eye-contact and the exposure of vulnerable body areas. Additionally, we predict that grooming modalities will vary according to kinship, sex, dominance rank, and age, reflecting adaptive behavioural strategies for managing social risk during social interactions.

## Materials and methods

### Study subjects

We studied two independent multimale–multifemale groups, one composed of rhesus macaques (*M. mulatta*) and the other of Tonkean macaques (*M. tonkeana*), over a 3-year period. Animals were housed at the Primate Center of the University of Strasbourg. The groups respectively consisted of 18 rhesus macaques (in 2021 and 2022) and 22 Tonkean macaques (in 2021 and 2023). The group composition is available in the supplementary (**Table S1**). Each group had free access to a 3100 m² and 3700m² wooded outdoor area and a 23m² and 19 m² heated indoor area equipped with structural enrichments (for the rhesus macaques and Tonkean macaques, respectively). Animals were fed with commercial monkey pellets once a day every day of the week, in addition to a supply of fresh fruit and vegetables once a week. Water was available *ad libitum*.

### Data collection

Data were collected directly in the wooded enclosure using a tablet (Samsung Galaxy Tab A), with the software ‘BORIS’ (Friard & Gamba, 2016) for Android over three years (March-July 2021, March-July 2022 for the rhesus macaques and March-August 2021, April-August 2023 for the Tonkean macaques). Data were recorded using 15-minute animal focal sampling in a pseudorandom order (each macaque was followed once before moving to the next sequence of focal). Each macaque was followed 15 times per year for a total of 135 hours for the rhesus macaques and a total of 168,75 hours for the Tonkean macaques. We recorded relative body orientation, and groomed body sites during focal sampling. ML collected the data for the Tonkean 2021 and Tonkean 2023. MJ collected the data for the rhesus 2021 (Kappa_MJxML_ = 0.95). NB collected 50% of the data of rhesus 2022 (Kappa_NBxML_ = 0.92). SS collected 50% of the data of rhesus 2022 (Kappa_SSxML_ = 0.91). Prior to the data collection phase, the kappas were calculated using five different animals during five focal samplings. Inter-observer reliability was assessed before data collection, with both observers recording the same focal samplings concurrently.

### Grooming modalities

The relative body orientation during grooming was divided into four categories: face-to-face, face-to-back, face-to-flank, and face-to-dorsal pelvis (**Figure 2A–D**). These four categories capture the full range of relative body orientations during grooming: face-to-face represents the highest degree of mutual frontal exposure, face-to-back and face-to-dorsal pelvis represent posterior exposure of the receiver, and face-to-flank represents lateral exposure. Together, they reflect a gradient of vulnerability and potential for eye-contact between partners. To determine the orientation category, we considered the emitter’s head orientation relative to the receiver’s body, regardless of the overall body posture of either individual. This coding rule was applied consistently across all postures, including cases where one or both individuals were lying down. Based on previous studies (Barton, 1985; Moser *et al.,* 1991; Pfoh *et al.,* 2021), we defined 11 possible body sites during grooming: face/ears, back head, back, limbs, flank, neck, chest, belly, ventral pelvis, dorsal pelvis, and armpit (**Figure 2E–F**). These two variables, relative body orientation and groomed body site were used to compute the Eye-Contact Probability score (ECP), described in the following section.

**Figure 2.**
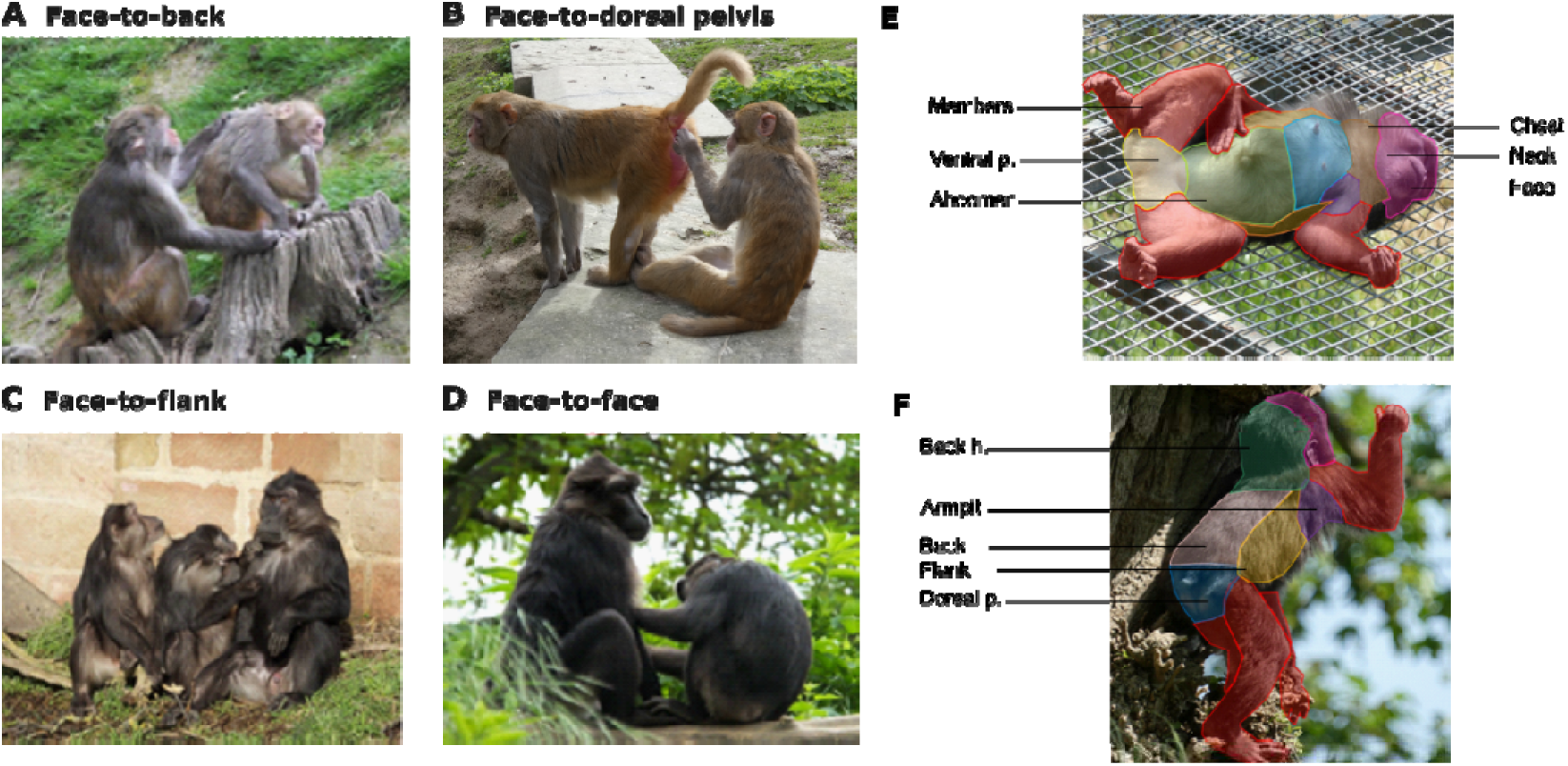
Grooming somatic modalities. **A–D**: Four relative body orientations during grooming: face-to-back (**A**), face-to-dorsal pelvis, (**B**), face-to-flank(**C**), and face-to-face (**D**). **E–F**: Body sites recorded during grooming. **E**: Ventral view (limbs: red; ventral pelvis: yellow; abdomen: green; chest: blue; neck: orange; face: purple). **F**: Dorsal view (back head: green; armpit: purple; back: grey; flank: yellow; dorsal pelvis: blue).

### Assessing Eye-Contact Probability (ECP)

As actual eye-contact between individuals is brief, rare and difficult to assess visually (but see Pedruzzi *et al.,* 2025), we did not quantify it directly but instead estimated the probability that eye-contact could occur during grooming. Using all-occurrences sampling, we filmed 541 allogrooming sessions, continuous grooming bouts between the same two individuals, without interruption or partner switch, comprising 391 sessions in rhesus macaques (≈3 h of grooming) and 150 sessions in Tonkean macaques (10–30 May 2023; ≈6 h). When several sessions overlapped, we filmed the one offering the best visibility, prioritising dyadic unidirectional grooming with identifiable individuals and visible eyes and groomed body site. Sessions were filmed until they ended or until video quality became insufficient, using a tripod-mounted camera or a camera trap. Relative body positions and groomed body parts were scored offline in BORIS (inter-observer reliability κ = 0.86, based on 64 min across four sessions).

To estimate ECP, we assessed how conducive each partner’s posture was to eye-contact, taking physiological constraints and each individual’s estimated visual field into account. Each partner received a score from 0 to 3 (0 = no orientation conducive to eye-contact; 1 = body oriented; 2 = body and face oriented; 3 = body, face and eyes oriented), the face counting only if the body was already oriented and the eyes only if the face was. Each configuration was assigned to a category *i* (0–6; Table S2) reflecting the combined effort both individuals would need to make eye-contact.

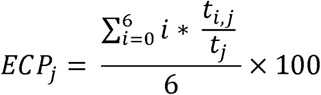

ECP_j_, ECP of the body part j

i, category from 0 to 6 assigned according to the orientation of each conspecific

t_i,j_, grooming time of the body part j with the score i

t_j_, total time of grooming of the body part j

The value of the ECP can range from 0 to 100 (maximum level of eye-contact probability)

For each body part *j*, ECP was computed from the grooming time spent in each category (*t*,) relative to the total grooming time of that part (*t*), and ranges from 0 (no eye-contact possible) to 100 (maximal eye-contact probability).

### Hierarchy of dominance

Dyadic agonistic interactions (including threats, supplants, chases, and physical assaults) and displacement interactions (approach-avoidance) were recorded ad libitum over the full study periods. Data were analysed using the ‘eloRating’ package on R (Neumann & Kulik, 2014) using David’s score (DS) (Gammell *et al*., 2003). In the analysis, we used the absolute difference in ordinal rank between the groomer and the individual being groomed (*rank difference*).

### Statistical analysis

#### Overview of the analytical approach

The analyses proceeded in two stages. First, we examined whether ECP varied across body positions and body parts using Wilcoxon rank-sum tests, in order to identify which grooming modalities were associated with higher or lower eye-contact probability. Second, we tested whether socio-demographic variables predicted the proportion of grooming time spent in positions or directed to body parts associated with high ECP, using a two-step GLMM approach.

### ECP comparisons

ECP values were compared across body positions and body parts using two-sample Wilcoxon rank-sum tests (Mann-Whitney U, exact procedure, R). Data from both species were analysed separately but pooled for the graphical representation, as the aim was to characterize general grooming modality patterns in terms of eye contact probability, irrespective of species differences. For body positions, face-to-face was compared separately to each of the three remaining positions (face-to-back, face-to-flank, and face-to-dorsal pelvis). For body parts, face, neck, and chest were compared to all remaining body parts combined. As individual identity was not systematically tracked across videos, some individuals may contribute multiple observations. Although this represents a limitation of the present study, the large and consistent effect sizes observed across all comparisons suggest that the overall pattern is robust. All tests were two-tailed and the significance threshold was set at p < 0.05.

### GLMM analyses

The objective of the present analyses was to determine which socio-demographic variables were associated with three aspects of grooming behaviour: (1) whether grooming occurred in the face-to-back position and, if so, the proportion of grooming time spent in that position (Pfb); (2) whether grooming occurred in the face-to-face position and, if so, the proportion of grooming time spent in that position (Pff); (3) whether grooming occurred on body parts associated with high eye-contact probability (ECP: face, chest, and neck) and, if so, the proportion of grooming directed to these body parts (Pecp). Face-to-face and face-to-back positions were retained for the main analyses as they represent the highest and lowest ECP values respectively (see Results, **Figure 3A**), and thus constitute the two extreme risk profiles in terms of eye-contact probability during grooming.

**Figure. 3.**
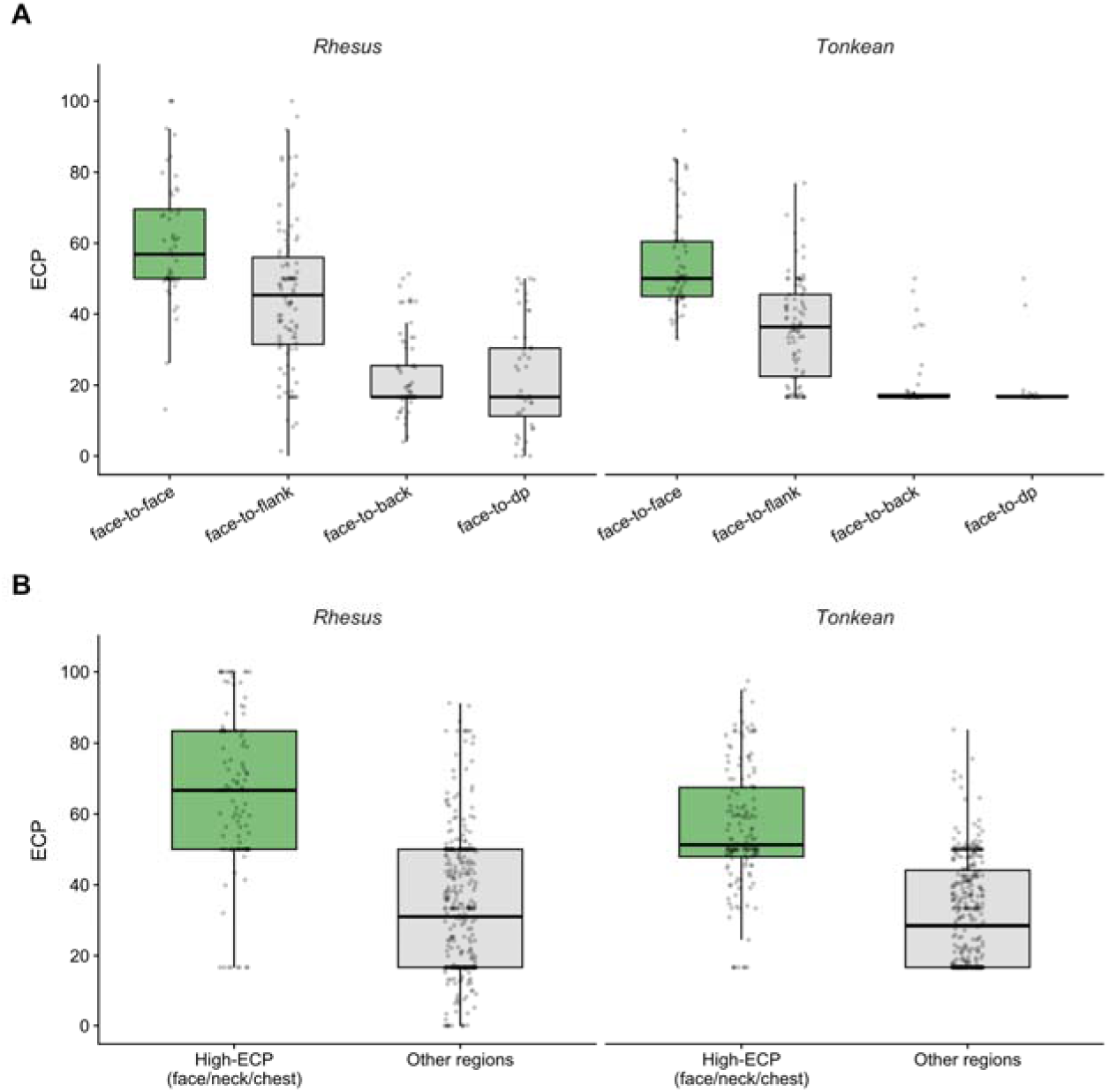
Eye-contact probability (ECP) during allogrooming in rhesus macaques and Tonkean macaques. **A** ECP across the four relative body positions; the face-to-face position (green) shows the highest values in both species. **B** ECP for the pooled high-ECP body parts (face, neck and chest; green) versus all other body parts combined (grey). Boxes show the median and interquartile range, whiskers extend to 1.5 × IQR, and individual grooming observations are overlaid as jittered points; n = number of grooming observations per category. In both species the grouped regions had a significantly higher ECP than the remaining regions (two-sided Wilcoxon rank-sum test: rhesus, W = 39,002, p < 0.001, r = 0.75; Tonkean, W = 49,601, p < 0.001, r = 0.70). *** p < 0.001

Each line of the dataset corresponded to a dyad. All possible dyads in each group were included, regardless of whether they groomed. For each of the three behavioural dimensions, analyses followed the same two-step structure (Liu *et al*., 2019), to accommodate the large number of dyads that never engaged in a given behaviour.

### Step 1: Presence/absence of the behaviour (binomial GLMM)

For the first step, a binomial generalized linear mixed model with a logit link was used to assess the influence of various predictors on the behaviour of interest (i.e., whether they ever groomed in a given position or targeted high-ECP body parts). Model diagnostics included checks for over dispersion and residual distribution using the *DHARMa* package. Multicollinearity among predictor variables was inspected using Variance Inflation Factors (VIF) computed with the *check_collinearity* function from the *performance* package (v0.12.0).

### Step 2: Proportion of grooming time (beta GLMM)

For dyads where the behaviour occurred (non-zero values), we modelled the proportion of grooming time using beta mixed-effects regression. Prior to fitting models, proportions were adjusted to avoid exact 0 or 1 values following standard recommendations (Baayen, 2008). Model significance was assessed using likelihood-ratio tests comparing the full model to reduced models excluding the term of interest. Model assumptions, including the absence of over dispersion, were verified using diagnostic functions from our custom scripts. Model stability was evaluated using *glmmTMB.stab* (source: glmmTMB_stability.r), and multicollinearity was again assessed using VIFs.

All analyses were conducted in R (R Core Team, 2024). Summary tables were generated using the *tab_model* function from the *sjPlot* package (v2.8.16). RV is the response variable, and the general formula used for all the model was:

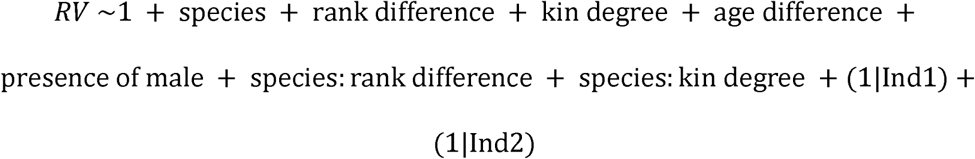

For all models, the predictors included species (RHESUS or TONKEAN), the absolute difference in rank between partners (rank difference), the absolute age difference (age difference), and the maternal kinship degree (kin degree: 0.5 for mother–offspring, 0.25 for siblings/grandparents, and 0.125 for aunt/uncle relationships; note that a kinship degree of 0.125 occurred only in the Tonkean macaque group). We also included male presence in the dyad (presence of male: TRUE if at least one male was present; FALSE otherwise), as males were hypothesized to increase risk during social interactions. Two interaction terms were specified: species × kinship degree, to model species differences in nepotism, and species × rank difference, to model species differences in dominance steepness. Both members of each dyad (Ind1 and Ind2) were included as random factors to account for the non-independence of dyadic observations, as each individual may appear in multiple dyads (Farine & Whitehead, 2015). Individual dyad members were arbitrarily assigned as Ind1 and Ind2 based on alphabetical order and changing this order does not affect the results of the models. In addition, to further verify that this assignment did not influence the results, all models were re-run with Ind1 and Ind2 swapped; fixed effect estimates were consistent across both assignments in terms of direction, magnitude, and significance. As each dyad contributes a single observation to the dataset, no dyad-level random effect was included. To facilitate interpretation of the parameter estimates, age difference, rank difference, and kinship degree were standardized (Schielzeth & Forstmeier, 2009) using z-transformation (scale function in R), setting mean = 0 and SD = 1. Wald tests were performed using the glmmTMB package.

## Results

### Level of Eye-Contact Probability Body positions

In both species, ECP was by far highest in the face-to-face position (rhesus: median = 56.86, IQR = 50.00–69.56, *n* = 53; Tonkean: median = 50.00, IQR = 45.00– 60.47, *n* = 64), which exceeded every other position (Wilcoxon rank-sum tests, all *p* < 0.001; rhesus: *W* ≥ 2,641; Tonkean: *W* ≥ 1,433; **Figure 3A**).

### Body parts

The face, neck and chest showed the highest ECP in both species (rhesus medians = 67.70, 60.44 and 70.05; Tonkean medians = 60.62, 50.47 and 49.61). When pooled, these three regions had a markedly higher ECP than all remaining body parts combined, in each species (rhesus: median = 66.67 vs 30.98; *W* = 39,002, *n□*= 116, *n□*= 385, *p* < 0.001, *r* = 0.75; Tonkean: median = 51.26 vs 28.52; *W* = 49,601, *n□*= 149, *n□*= 391, *p* < 0.001, *r* = 0.70; **Figure 3B**). Building on this result, we henceforth treat the face, neck and chest as a single category of high eye-contact body parts. In the analyses that follow, we no longer consider the ECP value itself, but instead use this category to test whether the two species differ in how much they groom these body parts.

### Face-to-back grooming

For the 151 dyads who groomed each other at least once in a face-to-back position, rhesus macaques and Tonkean macaques dedicated, on average 34.0 ± 3.0% and 56.6 ± 3.3% of their grooming time in the face-to back position, respectively. Rhesus macaques were more likely than Tonkean macaques to be involved at least once in a face-to-back grooming session (ß = -1.48, SE = 0.34, p < 0.001; **Table S3, Figure 4A**). Tonkean macaques allocated a smaller proportion of their grooming time to the face-to-back position than rhesus macaques (ß = -1.07, SE = 0.27, p < 0.001; **Table S3, Figure 4B**). Closely related dyads allocated a lower proportion of their grooming time to the face-to-back position than unrelated ones (ß = -0.43, SE = 0.09, p < 0.001) with a stronger effect in rhesus than in Tonkean macaques (ß = 0.42, SE = 0.14, p = 0.003; **Table S3, Figure 4C**).

**Figure 4.**
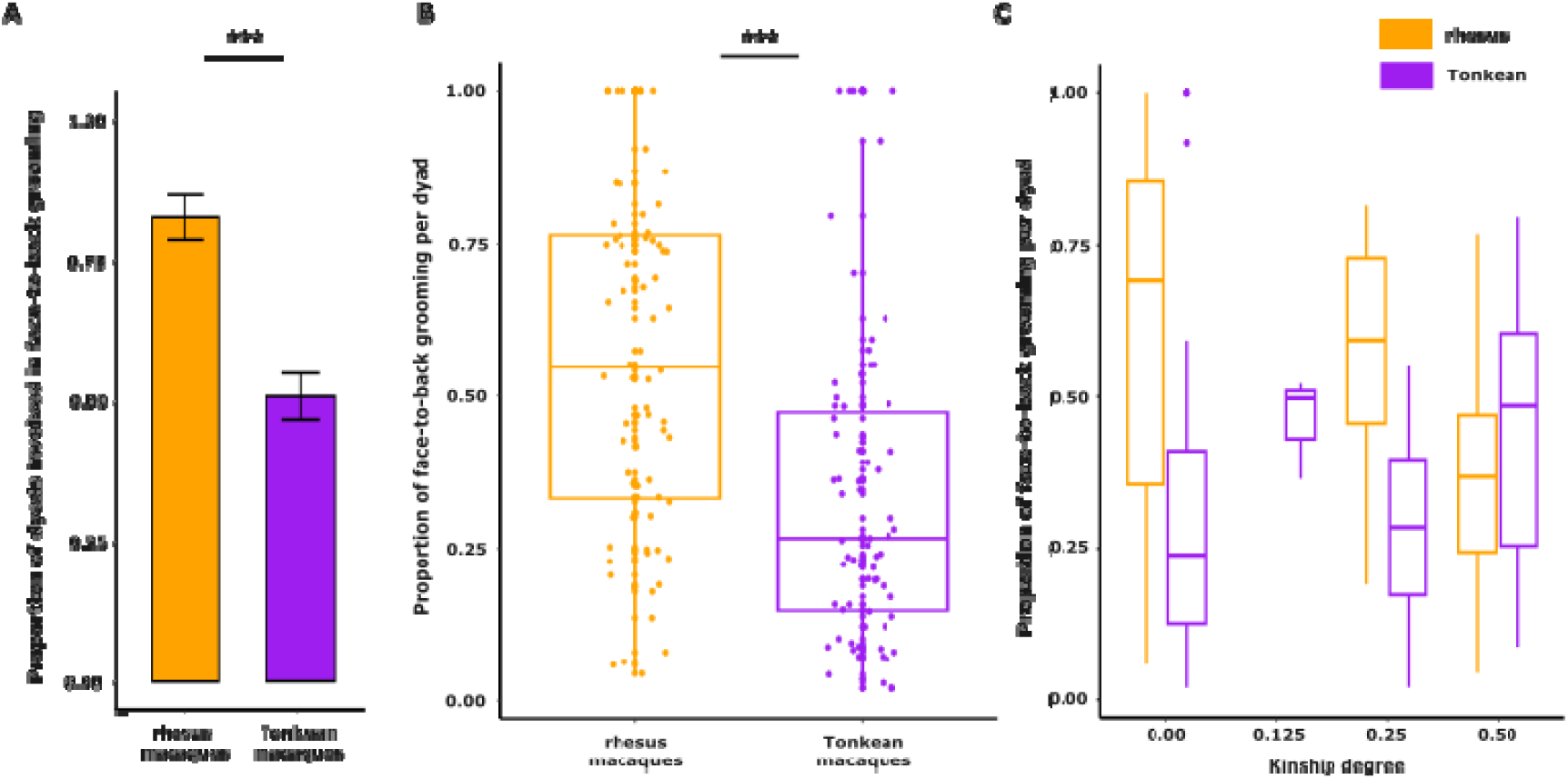
Effects of socio-demographic variables on the proportion of grooming time spent in the face-to-back position in rhesus and Tonkean macaques. Only significant predictors from the binomial GLMM (**A**) and beta GLMM (**B, C**) are shown. **A**: Proportion of dyads that engaged in face-to-back grooming in each species. **B**: Proportion of face-to-back grooming per dyad by species. **C**: Proportion of face-to-back grooming per dyad according to maternal kinship degree, shown separately for each species. ***p < 0.001 (Wald test). In all panels, rhesus and Tonkean macaques are depicted in orange and purple, respectively.

### Face-to-face grooming

For the 183 dyads who groomed each other at least once in a face-to-face position, Tonkean macaques and rhesus macaques dedicated, on average 58.4 ± 2.9 % and 47.8 ± 3.8 % of their grooming time in face-to-face position respectively. Tonkean macaques were more likely to be involved in face-to-face grooming sessions than rhesus macaques (ß = 0.81, SE = 0.34, p = 0.016; **Table S4, Figure S1A**). Closely related dyads were more likely to be involved in a face-to-face grooming session than other dyads (ß = 0.53, SE = 0.21, p = 0.009) with a stronger effect in rhesus macaque species (ß = -0.66, SE = 0.26, p = 0.010; **Table S4, Figure S1B**). The proportion of time allocated to this position was greater when at least one male was present in the dyad (ß = 0.49, SE = 0.20, p = 0.012; **Table S4**) than

### Grooming of high ECP body parts

For the 168 dyads who groomed at least once the high ECP body parts (face, chest, neck), Tonkean macaques and rhesus macaques dedicated, on average 33.8 ± 2.1 % and 22.3 ± 3.0 % of their grooming time on high ECP body parts. Tonkean macaques were more likely to groom high ECP body parts than rhesus macaques (ß = 0.93, SE = 0.36, p = 0.010; **Table S5**., **Figure 5A**). Dyads composed of at least one male were less likely to groom high ECP body parts than female-female dyads (ß = -0.87, SE = 0.34, p = 0.010; **Table S5**., **Figure 5B**). Closely related dyads were more likely to groom high ECP body parts than other dyads (ß = 0.38, SE = 0.17, p = 0.024), with a stronger effect in rhesus than in Tonkean macaques (ß =-0.56, SE = 0.22, p = 0.013; **Table S5**., **Figure 5C**). For the 168 dyads who groomed each other at least once those body parts, dyads with higher age difference were less likely to groom high ECP body parts (ß = -0.35, SE = 0.10, p < 0.001; **Table S5**., **Figure 5D**).

**Figure 5.**
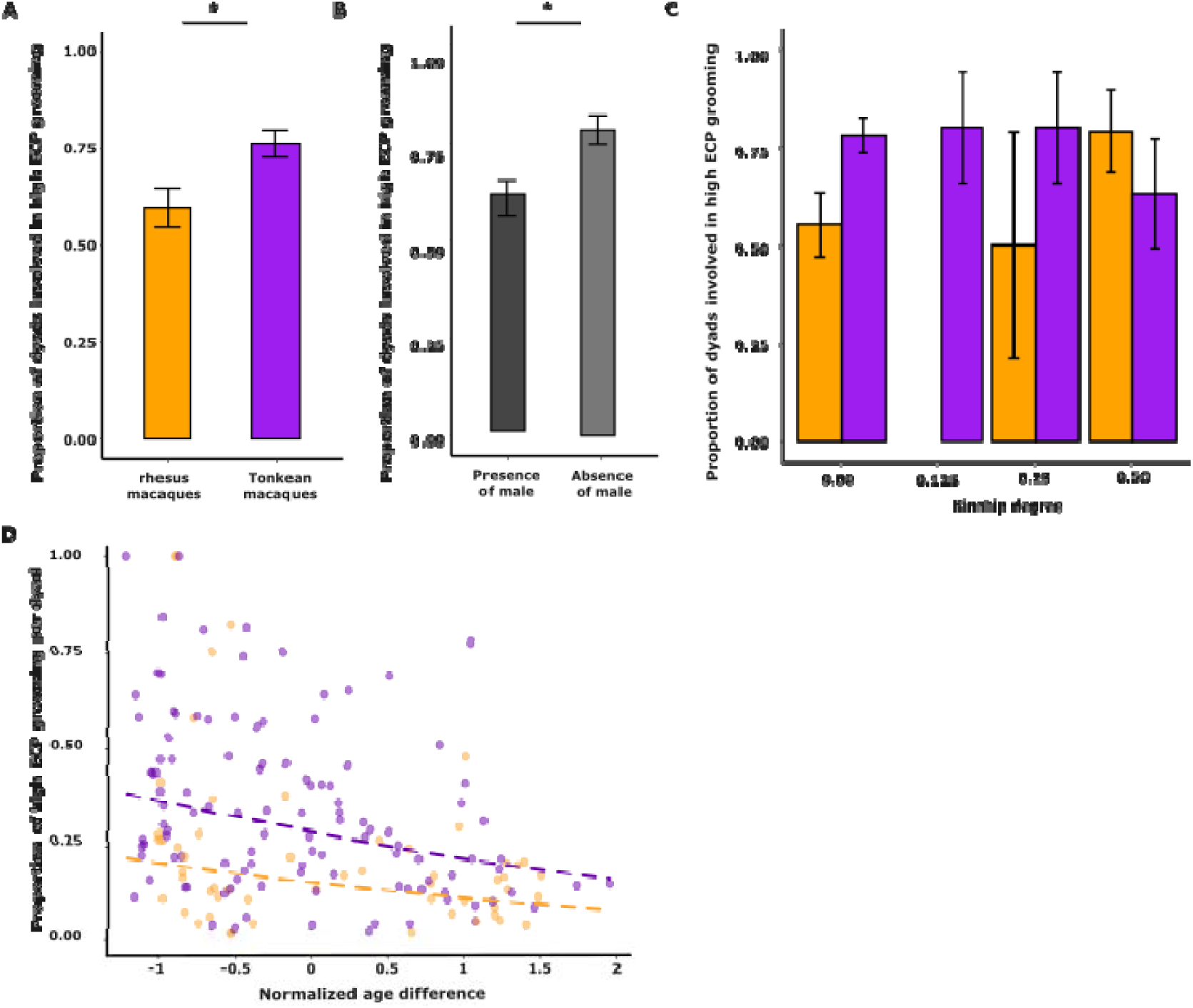
Effects of socio-demographic variables on the proportion of grooming time of high ECP body parts in rhesus and Tonkean macaques. Only significant predictors from the binomial GLMM (**A–C**) and beta GLMM (**D**) are shown. **A**: Proportion of dyads that engaged in high-ECP grooming in each species. **B**: Proportion of dyads showing high-ECP grooming according to the presence or absence of a male in the dyad. **C**: Proportion of dyads showing high-ECP grooming across maternal kinship degrees for each species. **D**: Proportion of grooming time allocated to high-ECP body parts per dyad. **\*** p < 0.05 (Wald test). In all panels, rhesus and Tonkean macaques are depicted in orange and purple, respectively.

## Discussion

The study of two macaque species differing in their level of social tolerance revealed that grooming modalities are associated with a combination of social and demographic factors, including kinship, age and male presence. These results suggest that risk management strategies during social interactions are modulated by species-typical social tolerance, with rhesus and Tonkean macaques showing distinct social grooming patterns.

We found that, compared to unrelated individuals, kin groomed each other in a face-to-face position. This kin-biased behavioural strategy also depended on species-specific social tolerance, as it was more pronounced in the more nepotistic rhesus macaques. Using our quantitative measure of Eye-Contact Probability (ECP), we further showed that related and same-age partners devoted more time to grooming body regions with high ECP (head, neck, chest). Overall, our results suggest that the effects of kinship on grooming modalities are consistent with the hypothesis that interactions between relatives are perceived as being less risky in NHPs. Indeed, kinship is generally associated with reduced risk: related females exhibit reduced aggression toward kin (Silk *et al*., 1981) and higher rates of reconciliation among relatives (Demaria & Thierry, 2001). Moreover, interacting with males may also involve elevated physical risk due to their greater potential for inflicting injury (Massey *et al*., 2022). While we found mixed results regarding grooming body orientation, our analysis revealed that the presence of male in a dyad significantly reduced high ECP grooming. Future work should also consider how interactions with reproductive behaviours may influence partner selection, grooming positions and which body sites are groomed (Colmenares, 1991).

Age and rank similarity may also reduce the likelihood of conflict escalation (de Waal & Luttrell, 1986), potentially enabling mutually beneficial interactions with lower anticipated costs. Dominance rank often plays a major role in structuring macaque societies, particularly in despotic species (Schino, 2001; King *et al*., 2011; Balasubramaniam *et al*., 2017). We therefore expected rank differences to influence grooming strategies, with interactions involving higher-ranking individuals carrying greater potential risk. However, in our dataset, dominance rank difference had no detectable effect on grooming strategies in either species. One possible explanation is that macaques adjust for rank-related risk *before* initiating grooming, namely during approach decisions thereby reducing the influence of rank once grooming has begun (Ballesta & Meunier, 2023). This interpretation aligns with previous work showing inconsistent or weak effects of hierarchy on social interaction dynamics (Nakamichi, 2003; Schino, 2001; Schino & Aureli, 2007). Although our study was not designed to quantify approach patterns directly, future analyses could examine whether dyads of similar versus dissimilar rank differ in their frequency of initiating grooming bouts, allowing a more explicit evaluation of pre-interaction filtering based on hierarchical position. It is also worth noting that rank difference alone may not fully capture the relevant variation in risk associated with dominance. Absolute rank position, independently of the distance between partners, could influence grooming strategies, as higher-ranking individuals may represent a source of risk regardless of the rank of their partner. Furthermore, kinship and rank are not independent in macaque societies, as maternal kin tend to occupy adjacent ranks within the hierarchy (Harcourt & Waal, 1992). This collinearity may have reduced our ability to detect independent effects of rank difference in the models. Future studies should consider disentangling these effects by including absolute rank as an additional predictor alongside rank difference, and by explicitly accounting for the overlap between kinship and rank proximity.

Species-typical social tolerance may also shape risk management strategies. We found that rhesus macaques (grade 1) relative to Tonkean macaques (grade 4) adopt more frequently face-to-back body orientation during grooming (34% and 56.6%, respectively). Similar patterns have been documented in other despotic species (Boccia, 1989), such as pig-tailed macaques (*M. nemestrina*, grade 2), which face away from groomer more often (51.3%) than the more tolerant bonnet macaques (*M. radiata*, grade 3; 22.3%). This strategy protects vulnerable body regions (Moser *et al*., 1991; Pfoh *et al*., 2021). In addition, our analysis shows that face-to-back orientation also reduces opportunities for eye-contact during grooming. Interestingly, similar interspecific differences in partner positioning have been documented in social play. (Petit *et al.,* 2008; Reinhart *et al.,* 2010) reported that more tolerant macaque species adopt face-to-face positions more frequently during play than despotic species, a pattern interpreted in terms of vulnerability and confidence between partners. The convergence of these findings across both grooming and play contexts suggests that body orientation as a risk management strategy may reflect a broader species-typical tendency shaped by social tolerance, rather than being context-specific.

Eye-contact carries an inherently ambivalent social valence in primates. On one hand, direct gaze can signal threat or aggression, and less tolerant species rely more heavily on gaze as a threat cue (Barton, 1985; Moser *et al*., 1991; Harrod *et al.,* 2020). Compared to other NHPs, macaques are generally considered relatively intolerant of direct eye-contact (Coss *et al.,* 2002). On the other hand, mutual gaze also supports bonding in both kin and non-kin relationships (Ehardt & Blount, 1984; Ferrari *et al*., 2009; Ballesta & Duhamel, 2015; Prinsen *et al.,* 2019), and has recently been shown to support shared engagement during affiliative interactions in macaques (Pedruzzi *et al.,* 2025). Specialized neural circuits integrate social and emotional cues during gaze processing (Emery, 2000; McFarland *et al.,* 2013; Mosher *et al.,* 2014; Gilardeau *et al*., 2021), suggesting that the valence attributed to eye-contact is not fixed but modulated by social context and relationship quality.

Against this background, our finding that Tonkean macaques groomed high-ECP regions more frequently than rhesus macaques is consistent with the idea that more tolerant species perceive eye-contact less negatively. In Tonkean macaques, the lower cost of conflict (Thierry, 2007; Thierry *et al.,* 1990) and the higher likelihood of reconciliation in tolerant species (Demaria & Thierry, 2001) may further reduce aversion to mutual gaze. Empirical evidence supports this: mutual gaze in macaques can exceed chance levels during face-to-face interactions, particularly in dyads with strong prosocial motivation (Ballesta & Duhamel, 2015).

To our knowledge, the specific conditions under which eye-contact during grooming triggers conflict escalation remain poorly documented. While gaze is established as a threat signal in agonistic contexts, its role in grooming-specific escalation has not been systematically examined. This represents an important gap, as understanding when eye-contact shifts from affiliative to threatening during grooming would considerably strengthen the risk management framework proposed here.

Overall, grooming positions that promote eye-contact may intentionally enhance affiliative engagement and signal trust, benefiting social cohesion. The valence attributed to eye-contact with particular partners can thus serve as a proxy for relationship quality, reflected in grooming dynamics. Our study therefore supports the proposal that grooming somatic modalities may serve as behavioural indicators of dyadic relationship quality and species-level social tolerance. The predominance of face-to-face grooming and high-ECP body sites in binomial rather than beta modelling indicate that the occurrence of such interactions may represents a signal of affiliation. Whether this signal is recognized and exploited by NHPs themselves remains an interesting avenue for future research.

In addition, our objective quantification of ECP allowed us to address a long-standing open question: whether individuals manage grooming strategies to protect vulnerable areas and/or prevent eye-contact exposure. Whereas earlier studies grouped all ventral body parts together as high eye-contact regions (Barton, 1985; Moser *et al*., 1991; Allanic *et al*., 2021; Pfoh *et al*., 2021), our analysis showed that only the head, neck, and chest exhibit high ECP, while the abdomen and ventral pelvis do not. Although the abdomen and ventral pelvis have low ECP, it can be argued that these two areas are particularly vulnerable, with the pelvis being a salient case in point. The results of this study thus align with the hypothesis that the management of eye-contact is a significant factor in determining the somatic modalities of grooming behaviour. It is important to note that the vulnerability-protection and eye-contact-avoidance hypotheses are not mutually exclusive and that further research is required to objectively assess body sites that are considered vulnerable, sensitive or even pleasant to be groomed in NHPs.

## Conclusion

Our findings indicate that macaques flexibly adjust their grooming strategies according to the socio-demographic identity of their partners and the species-typical degree of social tolerance. A central hypothesis emerging from our work is that more tolerant primate species perceive eye-contact less negatively. Tonkean macaques were more willing than rhesus macaques to groom high-ECP regions. Avoiding these regions may help prevent injury in higher-risk interactions, whereas engaging in face-to-face grooming may signal trust and deepen social bonds. Future studies could directly measure eye-contact valence using cognitive tasks (e.g. Deaner *et al*., 2005; Lio *et al*., 2019), include macaque species with intermediate tolerance grades, and quantify individual risk profiles (e.g., aggression frequency, injury rates). Taken together, these findings highlight that grooming somatic modalities are information-rich, as macaques appear to adjust them to manage social risk and to express relationship quality

## Data Availability Statement

The data that support the findings of this study are openly available on github: https://github.com/MLprimatology/Risk-management-strategies-in-social-grooming-among-Tonkean-and-rhesus-macaques.git

## Supporting information

Supplementary informations

## Acknowledgement

The authors thank the University of Strasbourg, the CNRS and Silabe (https://www.silabe.com/) for supporting this research and providing animal care. The work was funded by ANR-21-CE37-0016 to S.B. and H.M.

## Statement of Ethics

This non-invasive research complied with French legal requirements for the use of animals in research and EU Directive 2010/63/EU on the welfare of animals used for scientific purposes. As the study was purely observational, no ethical approval was required.

