## Supplementary informations for "Risk management strategies in social grooming among Tonkean and rhesus macaques"

**Macaques group composition**

**Table S1**. Detailed information of the social groups at the time of the study. The name with the three first letters of the name, sex (F for female and M for male), Kinship: matrilineal lineage identifier. Individuals sharing the same number belong to the same matriline, age (at the beginning of the focal sampling in year) and the rank (ordinal rank in the dominance hierarchy for each year). If empty, cases correspond to absent or too young subjects. **A**. for the Tonkean macaques in 2021 and 2023 and **B**. for the group of rhesus macaques in 2021 and 2022.

A. **Tonkean macaques	B.  rhesus macaques**

|  | | | **2021** | | **2023** | |
| --- | --- | --- | --- | --- | --- | --- |
| **Name** | ***Sex*** | ***Kinship*** | ***Age*** | ***Rank*** | ***Age*** | ***Rank*** |
| abr | M | 3 | 7.93 | 4 |  |  |
| ala | M | 4 | 8.08 | 3 |  |  |
| bar | M | 4 | 6.37 | 7 |  |  |
| ber | F | 1 | 6.43 | 12 | 8.47 | 5 |
| ces | M | 4 | 5.87 | 18 |  |  |
| dor | F | 2 | 5.07 | 20 | 7.11 | 13 |
| eri | M | 1 | 4.05 | 21 | 6.10 | 8 |
| fic | F | 2 | 3.15 | 22 | 5.19 | 6 |
| gai | F | 1 | 1.48 | 17 | 3.52 | 18 |
| gan | M | 2 | 2.00 | 16 | 4.04 | 20 |
| han | F | 4 |  |  | 2.67 | 21 |
| hav | F | 3 |  |  | 3.10 | 16 |
| her | M | 4 |  |  | 2.65 | 17 |
| hor | M | 2 |  |  | 2.67 | 14 |
| ind | M | 1 |  |  | 1.50 | 22 |
| iro | M | 2 |  |  | 1.33 | 19 |
| isi | F | 1 |  |  | 1.90 | 15 |
| jea | F | 5 | 25.36 | 8 | 27.40 | 10 |
| lad | F | 1 | 23.53 | 14 |  |  |
| las | F | 1 | 10.04 | 19 | 12.08 | 12 |
| nem | F | 2 | 10.28 | 11 | 12.33 | 3 |
| ner | F | 2 | 22.27 | 5 | 24.31 | 2 |
| olg | F | 3 | 20.65 | 9 | 22.70 | 7 |
| oll | M | 3 | 10.39 | 2 |  |  |
| pac | M | 4 | 9.76 | 1 | 11.80 | 1 |
| pat | F | 4 | 19.65 | 10 | 21.69 | 11 |
| uly | M | 5 | 16.25 | 6 |  |  |
| yin | F | 2 | 11.77 | 13 | 13.81 | 4 |
| yoh | F | 4 | 11.74 | 15 | 13.78 | 9 |

|  | | | **2021** | | **2022** | |
| --- | --- | --- | --- | --- | --- | --- |
| **Name** | ***Sex*** | ***Kinship*** | ***Age*** | ***Rank*** | ***Age*** | ***Rank*** |
| any | F | 7 | 0.82 | 14 | 1.91 | 15 |
| arw | F | 1 | 2.67 | 17 | 3.75 | 4 |
| baa | M | 7 |  |  | 0.96 | 17 |
| bor | M | 2 | 2.86 | 7 | 3.94 | 3 |
| djo | M | 2 | 1.85 | 11 | 2.93 | 6 |
| eow | F | 5 | 2.85 | 16 | 3.93 | 12 |
| far | M | 3 | 2.82 | 15 | 3.90 | 7 |
| kab | F | 2 | 21.74 | 3 | 22.82 | 2 |
| lin | F | 7 | 13.05 | 4 |  |  |
| nat | F | 4 | 0.58 | 13 | 1.66 | 16 |
| qui | F | 1 | 21.91 | 6 | 22.99 | 8 |
| sam | M | 3 |  |  | 0.93 | 18 |
| spl | M | 7 | 1.83 | 12 | 2.91 | 10 |
| the | M | 4 | 2.77 | 5 | 3.85 | 11 |
| uli | M | 6 | 18.90 | 1 | 19.98 | 1 |
| veg | F | 4 | 7.85 | 2 |  |  |
| vla | M | 3 | 0.94 | 18 | 2.02 | 13 |
| vol | F | 3 | 17.88 | 8 | 18.96 | 5 |
| yel | F | 1 | 0.84 | 10 | 1.95 | 9 |
| yva | M | 2 | 0.53 | 9 | 1.61 | 14 |

**ECP Measurement**

**Table S2**: Possible orientations of the emitter (E) and the receiver (R) during social grooming. We considered whether individuals had to move their eyes, their head and/or their full body to make eye-contact with their partner, based on individuals estimated visual field. We adopted a score from 0 to 3 for both the emitter and the receiver of the grooming. We scored 0 when neither the body, nor the face, nor the eyes were conducive to eye-contact. We scored 1 when only the body was conducive to eye-contact, but not the face or eyes. We scored 2 when the body and face were conducive to eye-contact, but not the eyes, and we scored 3 when the body, face and eyes were conducive to eye-contact. Note that we considered that the face could be conducive to eye-contact only if the body was. Therefore, E0 means that the body of the emitter isn’t conducive to eye-contact (R0 the receiver); E1 only the body of the emitter is conducive to eye-contact (R1 for the receiver). E2 means that the body and face of the emitter are conducive to eye-contact (R2 for the receiver). E3: body, face and eyes are conducive to eye-contacts (R3 for the receiver). Similarly, the eyes could be conducive to eye-contact only if the face was too. We assigned each possible situation (e.g. E0R1, E2R2) to a category, according to the efforts needed to make eye-contact. From *i=*0 (both bodies must move) to *i*=6 (if only up to one individual has to move his eyes to make eye-contact).

| **Notation (Emitter Receiver)** | **i category** |
| --- | --- |
| E3R3, E2R3, E3R2 | 6 |
| E2R2 | 5 |
| E1R3, E3R1 | 4 |
| E1R2, E2R1 | 3 |
| E1R1 | 2 |
| E3R0, E2R0, E1R0, E0R1, E0R2, E0R3 | 1 |
| E0R0 | 0 |

**Face-to-back grooming**

**Table S3**: Results of the two-step GLMMs testing how socio-demographic factors (maternal kin degree of the dyads, rank and age difference of the conspecifics of the dyad, the presence of male and the species) influence the involvement or not in a face-to-back grooming session (fb) and the proportion of face-to-back grooming among the grooming time of the dyads involved in a grooming session. Estimates and Standard Error (SE) are presented in the table. Variables with p-value inferior to 0.05 (in bold) had a significant effect on the response variables (Wald test). The columns on the left correspond to the model testing for the presence or absence of behavior (Step 1), while the columns on the right correspond to analyzing the proportion of behavior observed (Step 2).

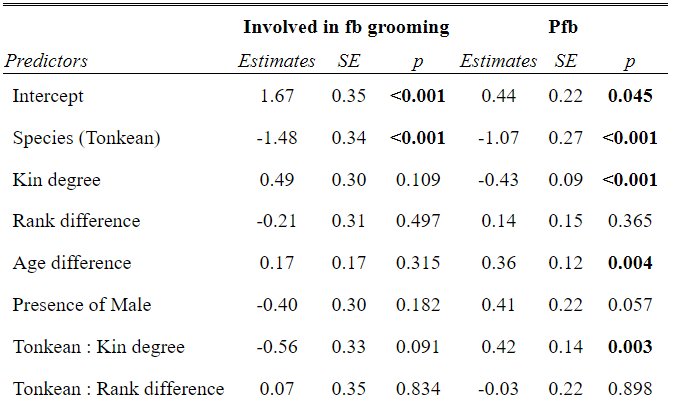

**Face-to-face grooming**

**Table S4**: Results of the two-step GLMMs testing how socio-demographic factors (maternal kin degree of the dyads, rank and age difference of the conspecifics of the dyad, the presence of male and the species) influence the involvement or not in a face-to-face (ff) grooming session and the proportion of face-to-face (Pff) grooming among the grooming time of the dyads involved in a grooming session. Estimates and Standard Error (SE) are presented in the table. Variables with p-value inferior to 0.05 (in bold) had a significant effect on the response variables (Wald test). The columns on the left correspond to the model testing for the presence or absence of behavior (Step 1), while the columns on the right correspond to analyzing the proportion of behavior observed (Step 2).

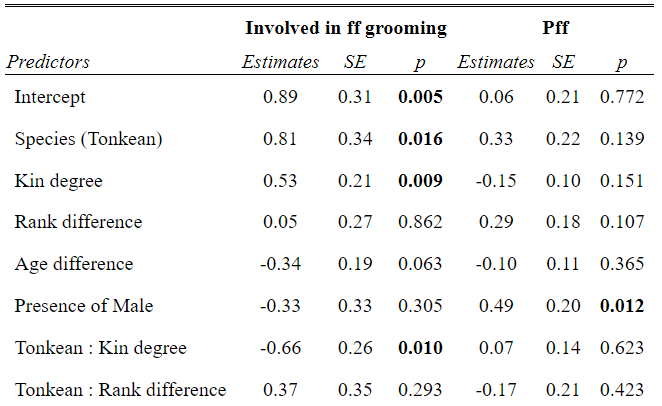

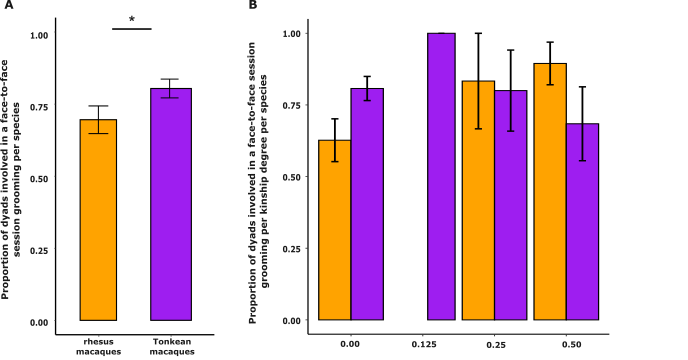

**Figure S1.** Influence of socio-demographic variables on the proportion of dyads involved in a face-to-face position during grooming in rhesus and Tonkean macaques. Only significant predictors from the binomial GLMM are shown. (A) Proportion of dyads engaging in at least one face-to-face grooming session in each species, with rhesus macaques in orange and Tonkean macaques in purple. (B) Proportion of dyads engaging in at least one face-to-face grooming session according to maternal kinship degree and species, with rhesus macaques in orange and Tonkean macaques in purple. * p < 0.05 (Wald test).

**Grooming of high ECP body parts**

**Table S5**: Results of the two-step GLMMs testing how socio-demographic factors (maternal kin degree of the dyads, rank and age difference of the conspecifics of the dyad, the presence of male and the species) influence the involvement or not in a high ECP grooming session and proportion of high ECP grooming (Pecp) among the grooming time of the dyads involved in a grooming session. Estimates and Standard Error (SE) are presented in the table. Variables with p-value inferior to 0.05 (in bold) had a significant effect on the response variables (Wald test). The columns on the left correspond to the model testing for the presence or absence of behavior (Step 1), while the columns on the right correspond to analyzing the proportion of behavior observed (Step 2).

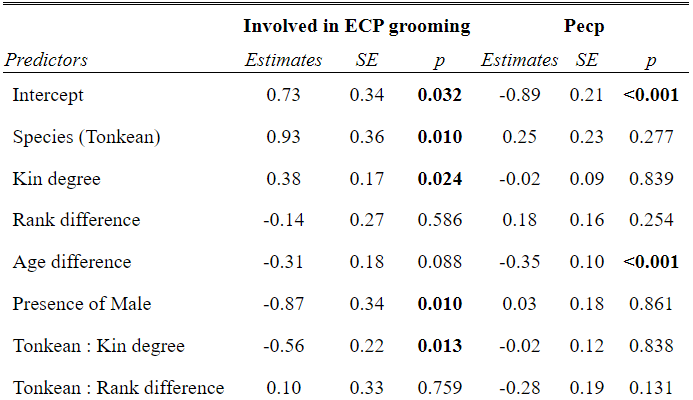
